# Preeclampsia-associated piRNA Regulates Trophoblast Function via YTHDF2-Mediated m6A Methylation

**DOI:** 10.1101/2025.04.07.647678

**Authors:** Ying Zhao, Shuang Liang, Miao Guo, Shaoyuan Huang, Houzhi Yang, Xu Chen, Shan-Shan Li, Xin Jin

**Author notes:** Corresponding Author Xin Jin School of Medicine, Nankai University, Tianjin, China Shan-Shan Li School of Medicine, Nankai University, Tianjin, China. Xu Chen Tianjin Central Hospital of Gynecology Obstetrics, Tianjin, China. These authors contributed equally to this study (Ying Zhao, Shuang Liang).

## Abstract

Preeclampsia is marked by abnormal placental development and impaired trophoblast function. We identified Preeclampsia-associated trophoblast piRNA (PETPIR) as an important regulator of trophoblast function and m6A modifications. Placental samples from preeclamptic and healthy pregnancies were analyzed using piRNA microarrays and m6A epitranscriptomic profiling. Functional assays in trophoblast cells, transcriptomic analysis, and an in vivo PETPIR-overexpressing mouse model were employed to investigate PETPIR’s mechanistic roles. We found that PETPIR was significantly upregulated in preeclamptic placentas. PETPIR inhibited trophoblast cell proliferation, migration, and invasion by inducing cell cycle arrest and apoptosis. Transcriptome analysis demonstrated that PETPIR activated hypoxia-responsive pathways. Additionally, PETPIR enhanced global m6A methylation and stabilized NDRG1 transcripts by interacting with the m6A reader YTHDF2. In vivo, PETPIR overexpression in pregnant mice recapitulated key features of preeclampsia, including maternal hypertension and fetal growth restriction. These results suggest PETPIR is a key player in the pathogenesis of preeclampsia, functioning through the PETPIR/YTHDF2/NDRG1 axis to drive hypoxia and m6A epitranscriptomic dysregulation. We provide novel insights into preeclampsia’s molecular mechanisms, highlighting piRNA as a promising biomarker and therapeutic target.

## Introduction

Preeclampsia is a life-threatening hypertensive disorder of pregnancy that significantly contributes to maternal and fetal morbidity and mortality (1, 2). Diagnosed after 20 weeks of gestation, it is characterized by the sudden onset of hypertension and at least one associated complication, such as proteinuria, maternal organ dysfunction, or uteroplacental dysfunction (3, 4). Despite its clinical and public health significance, the etiology of preeclampsia remains inadequately defined, particularly in its term and postpartum forms (5). This lack of understanding poses a major obstacle to the development of effective predictive tools and targeted therapeutic interventions.

The core of preeclampsia’s pathogenesis is placental dysfunction, driven by impaired trophoblast invasion and inadequate spiral artery remodeling (6). These defects result in placental hypoperfusion and chronic hypoxia, which initiate a cascade of systemic effects, including oxidative stress, inflammation, and widespread endothelial dysfunction (7). Recent evidence highlights the importance of non-coding RNAs (ncRNAs), including microRNAs (miRNAs), circular RNAs (circRNAs), and long non-coding RNAs (lncRNAs), in regulating key aspects of trophoblast biology and placental function (8–12). These ncRNAs act as critical regulators of gene expression and epigenetic modifications, regulating processes such as angiogenesis, immune responses, and hypoxia adaptation. However, their specific roles in trophoblast dysfunction and the pathophysiology of preeclampsia remain underexplored.

PIWI-interacting RNAs (piRNAs) are a distinct class of small ncRNAs that have recently emerged as important regulators of gene expression beyond their traditional role in transposon silencing in germ cells (13, 14). piRNAs are derived from long single-stranded precursor transcripts. These precursors undergo a complex maturation process, including fragmentation, trimming, and 3′-end methylation, resulting in mature piRNAs that bind to PIWI proteins (15, 16). While piRNAs are essential for maintaining genomic integrity in germ cells, growing evidence has expanded their function to somatic tissues. In these contexts, piRNAs are implicated in a variety of cellular processes, including epigenetic modifications, RNA stability, and protein translation (17). Their versatile roles have garnered significant attention for their potential involvement in human diseases (18, 19). Despite these advances, the role of piRNAs in placental biology and pregnancy-related disorders such as preeclampsia remains largely unexplored. Recent studies have revealed that piRNAs influence the epitranscriptomic landscape, including the regulation of N6-methyladenosine (m6A) modifications—a prevalent RNA modification that modulates RNA metabolism, stability, and translation efficiency. Emerging evidence suggests that piRNAs may act as upstream regulators of m6A methylation machinery, linking piRNA-mediated m6A regulation to the pathogenesis of cancers (20–24). This highlights the functional versatility of piRNAs as m6A epigenetic effectors.

The significance of m6A modifications extends to pregnancy-related disorders, particularly preeclampsia, where dysregulated m6A patterns have been implicated in placental dysfunction and trophoblast abnormalities (25–27). These pathological features are accompanied by changes in RNA methylation, suggesting that m6A is pivotal in shaping the molecular underlying of preeclampsia (28). Despite growing recognition of m6A’s importance in placental biology, the regulatory interplay between piRNAs and m6A remains unexplored. This raises critical questions about whether piRNAs contribute to the m6A reprogramming observed in preeclampsia. Elucidating the molecular mechanisms by which piRNAs influence m6A modifications in trophoblasts could provide novel insights into their roles in placental development and the pathogenesis of preeclampsia, offering promising avenues for targeted diagnostics and therapeutics.

In this study, we identify a novel piRNA, termed Preeclampsia-associated trophoblast piRNA (PETPIR), significantly upregulated in preeclamptic placentas. We explore its functional role in trophoblast biology and its regulatory mechanisms. Our findings demonstrate that PETPIR modulates trophoblast function through YTHDF2-mediated m6A methylation. By stabilizing hypoxia-responsive transcripts such as NDRG1, PETPIR contributes to the hypoxic conditions characteristic of preeclampsia. Using a PETPIR-overexpressing mouse model, we recapitulate key features of the disease, including hypertension and fetal growth restriction, establishing PETPIR as a functional driver of preeclampsia. This study provides novel insights into the molecular mechanisms of preeclampsia, highlighting the role of piRNAs in epitranscriptomic regulation and trophoblast function. Identifying PETPIR as a key player in preeclampsia pathophysiology offers a promising avenue for future research into its potential as a biomarker and therapeutic target.

## Methods

### Patients and samples

This study involved the collection of placental tissues and plasma samples from pregnant women recruited at Tianjin Central Hospital of Obstetrics and Gynecology between October 2021 and December 2022 (Clinical Trial Registration Number: ChiCTR2100043520; Ethics Approval Number: 2021KY042). The study protocol adhered to the guidelines of the ethics committee of Tianjin Central Hospital of Obstetrics and Gynecology. Written informed consent was obtained from all participants before sample collection. Participants included pregnant women diagnosed with preeclampsia (PE) and healthy controls. The diagnosis of PE was based on the criteria established by the American College of Obstetricians and Gynecologists (ACOG). PE was defined as a systolic blood pressure of ≥ 140 mmHg or diastolic blood pressure of ≥ 90 mmHg on two separate occasions at least 4 hours apart after 20 weeks of gestation in women with previously normal blood pressure, accompanied by significant proteinuria (≥300 mg protein per 24-hour urine collection, a protein/creatinine ratio of ≥ 0.3 mg/dL, or a dipstick reading of 2+ in a random urine sample). Detailed clinical characteristics of the participants are summarized in Table 1. Placental tissue and plasma samples were collected at the time of cesarean delivery. Placental tissues were snap-frozen in liquid nitrogen and stored at −80 °C. Venous blood samples were collected, and plasma was separated by centrifugation at 3000 rpm for 10 minutes at 4 °C. The supernatants were stored at −80 °C until further analysis.

**Table 1.** Clinical characteristics of the pregnant women enrolled in this study.

|  | Control (n=29) | PE (n=29) |
| --- | --- | --- |
| Maternal age, years | 32.1 ± 4.6 | 31.3 ± 3.1 |
| Body-Mass Index, kg/m<sup>2</sup> | 26.7 ± 4.2 | 28.9 ± 6.1 |
| Gestational Age, days | 248.8 ± 19.7 | 246.4 ± 21.1 |
| Systolic Blood Pressure, mmHg | 110.8 ± 7.7 | 162.9 ± 17.4 * |
| Diastolic Blood Pressure, mmHg | 67.9 ± 5.6 | 101.1 ± 11.5 * |
| Urine Protein (g/24h) | - | 2.8 ± 2.0* |
| Infant Birth Weight (g) | 2553 ± 734 | 2087 ± 663* |

Values are mean ± SD; P values are only shown for the significant differences between groups.
\* P<0.05.

### piRNA Microarray Analysis

RNA quantity and integrity were assessed using a NanoDrop ND-1000 Spectrophotometer (Thermo Scientific) and denaturing agarose gel electrophoresis. piRNA profiling was performed using the Arraystar HG19 piRNA Array (Aksomics Inc), with piRNAs mapped to the HG19 genome sequence from the NCBI database using UCSC Blat. The array included approximately 23,000 piRNAs with high-quality genomic matches, designed via a duplex method. For labeling, 1 µg of total RNA was 3′-end-labeled with a Cy3 fluorescent dye using T4 RNA ligase (Exiqon). Cy3-labeled RNA was hybridized to the array in Agilent SureHyb Hybridization Chambers at 65°C for 17 hours. After hybridization, slides were washed in an ozone-free environment and scanned with the Agilent DNA Microarray Scanner (G2505C). Data were analyzed using Agilent Feature Extraction software (v11.0.1.1) and normalized with the GeneSpring GX software package (v12.1, Agilent Technologies). Differentially expressed piRNAs were identified using fold change filtering.

### Cell Culture and Transfection

The human extravillous trophoblast cell line HTR-8/SVneo (Zhongqiao Xinzhou Biotechnology, Shanghai, China) was cultured in RPMI-1640 medium supplemented with 10% fetal bovine serum (FBS) and 1% penicillin-streptomycin (Gibco, Thermo Fisher Scientific) at 37°C in a humidified incubator with 5% CO₂. For transfection, a piRNA-1040 mimic and negative control (NC) were synthesized by GenePharma (Shanghai, China). Transfections were performed using Lipofectamine 2000 (Invitrogen, USA) according to the manufacturer’s protocol. Briefly, HTR-8/SVneo cells were seeded in 6-well plates to 70–80% confluency. The piRNA-1040 mimic or NC was diluted in Opti-MEM (Gibco) and mixed with Lipofectamine 2000. The transfection complexes were added to cells and incubated for 6 hours, followed by medium replacement with RPMI-1640 containing 10% FBS. Cells were harvested 48 hours post-transfection for downstream analyses.

### RNA Fluorescence In Situ Hybridization

A Cy3-labeled piRNA-1040 probe was synthesized by GenePharma (Shanghai, China). HTR-8/SVneo cells were cultured on glass coverslips, fixed with 4% paraformaldehyde for 15 minutes at room temperature, and permeabilized with 0.5% Triton X-100 in PBS for 10 minutes. Hybridization was carried out overnight at 37°C in a humidified chamber using the Cy3-labeled probe. Post-hybridization, cells were washed with saline-sodium citrate (SSC) buffer to remove unbound probes. Nuclei were counterstained with DAPI, and coverslips were mounted with an anti-fade mounting medium. Fluorescence images were captured using a confocal microscope (STELLARIS, Leica, Germany).

### Nuclear and Cytoplasmic RNA Extraction

Nuclear and cytoplasmic RNA fractions were isolated using the PARIS™ Kit (Thermo Fisher Scientific, AM1921) according to the manufacturer’s protocol. HTR-8/SVneo cells were lysed in a cell fractionation buffer, and nuclear and cytoplasmic fractions were separated by centrifugation. RNA was extracted from each fraction, and its quality and quantity were measured using a NanoDrop spectrophotometer (Thermo Fisher Scientific). Quantitative PCR (qPCR) was performed to assess RNA abundance in the fractions, with U6 small nuclear RNA and ACTIN mRNA serving as controls for nuclear and cytoplasmic localization, respectively.

### Cell Proliferation, Cell Cycle, and Apoptosis Assays

Cell proliferation was evaluated using the Cell Counting Kit-8 (CCK-8) assay (Beyotime Biotechnology, Shanghai, China). HTR-8/SVneo cells (2 × 10³ cells/well) were seeded in 96-well plates. After adherence, 10 μl of CCK-8 reagent was added to each well, and the plates were incubated at 37°C for 1 hour. Absorbance at 450 nm was measured using a microplate reader to determine cell viability.

Cell cycle distribution was assessed using the Cell Cycle Analysis Kit (Beyotime Biotechnology, Shanghai, China). HTR-8/SVneo cells were fixed in 70% ethanol overnight at 4°C, washed with phosphate-buffered saline (PBS), and stained with propidium iodide (PI) solution containing RNase A. Samples were analyzed by flow cytometry (BD Biosciences, USA) to quantify the proportions of cells in G0/G1, S, and G2/M phases.

Apoptosis was detected using the Annexin V-FITC/PI Apoptosis Detection Kit (Solarbio, Beijing, China). HTR-8/SVneo cells were resuspended in 1× binding buffer, stained with Annexin V-FITC and PI, and incubated in the dark at room temperature for 15 minutes. Stained cells were analyzed immediately by flow cytometry to quantify early and late apoptotic populations.

### Wound Healing Assay

The migratory capacity of HTR-8/SVneo cells was assessed using a wound healing assay. Cells were seeded in 6-well plates and grown to near confluence. After transfection with the respective constructs, a straight wound was created in the monolayer using a sterile pipette tip. Detached cells were removed by washing with phosphate-buffered saline (PBS), and a fresh culture medium was added. Images of the wound area were captured immediately after scratching (0 h) using an inverted microscope. The plates were incubated at 37°C with 5% CO₂, and the wound area was re-imaged after 24 hours. Migration rates were calculated based on the percentage reduction in the wound area using ImageJ software.

### Transwell Assay

Cell invasion and migration were evaluated using transwell chambers (Corning, USA) with or without Matrigel coating. For the invasion assay, transwell inserts were pre-coated with Matrigel and incubated at 37°C for 1 hour to allow polymerization. HTR-8/SVneo cells were transfected, incubated for 24 hours, harvested, and resuspended in serum-free RPMI-1640 medium. A total of 200 μL of the cell suspension (approximately 2 × 10⁴ cells) was added to the upper chamber, while 600 μL of RPMI-1640 containing 10% fetal bovine serum (FBS) was placed in the lower chamber as a chemoattractant. Cells were incubated at 37°C with 5% CO₂ for 48 hours. Non-invading cells on the upper membrane surface were removed with a cotton swab. Invaded cells on the lower surface were fixed with 4% paraformaldehyde for 15 minutes and stained with 0.1% crystal violet (Solarbio, Beijing, China) for 20 minutes. Stained cells were imaged under a light microscope, and cell counts were obtained from three randomly selected fields per membrane. Migration assays followed the same protocol but without Matrigel coating on the transwell inserts.

### RNA Sequencing Analysis

Total RNA was extracted from HTR-8/SVneo cells using TRIzol Reagent (Invitrogen, USA) according to the manufacturer’s instructions. RNA purity was measured with a kaiaoK5500® Spectrophotometer (Kaiao, Beijing, China), and RNA integrity and concentration were evaluated using the RNA Nano 6000 Assay Kit and Bioanalyzer 2100 system (Agilent Technologies, USA). For RNA-seq library preparation, 3 μg of high-quality RNA per sample was processed using the NEBNext Ultra RNA Library Prep Kit for Illumina (NEB, USA) following the manufacturer’s protocol. Libraries were sequenced on the Illumina NovaSeq 6000 platform (Shanghai Personal Biotechnology Co., Ltd.). Quality control of raw reads was performed with FastQC, and adapter sequences and low-quality bases were removed using Cutadapt. Clean reads were aligned to the human reference genome (hg38) using HISAT2. Transcript assembly for each sample was carried out with StringTie, and transcriptomes were merged across all samples using gffcompare to generate a unified annotation. Transcript abundance was quantified as Fragments Per Kilobase of transcript per Million mapped reads (FPKM). Differentially expressed genes (DEGs) were identified using the R package DESeq2, with significance thresholds set at an adjusted p-value < 0.05 and |log2(fold change)| > 1.

### m6A Dot Blot Assay

Total RNA was extracted using TRIzol Reagent (Invitrogen, Thermo Fisher Scientific) following the manufacturer’s protocol. Purified RNA was denatured at 95°C for 3 minutes and immediately cooled on ice. Equal volumes of denatured RNA were spotted onto Amersham Hybond-N+ membranes (Solarbio, YA1760) and UV crosslinked at 254 nm with 0.12 J/cm² to immobilize the RNA. Membranes were blocked with 5% non-fat milk in TBST (Tris-buffered saline with 0.1% Tween-20) for 1 hour at room temperature to prevent non-specific binding. They were then incubated overnight at 4°C with an anti-m6A antibody (1:5000, Abcam, ab314476). After TBST washes, membranes were incubated with an HRP-conjugated secondary antibody (1:5000) for 1 hour at room temperature. Visualization was performed using an ECL chemiluminescence detection system (Thermo Fisher Scientific). To confirm equal RNA loading, membranes were stained with 0.02% methylene blue for 10 minutes and rinsed with RNase-free water until RNA dots were visible. m6A signal intensity was quantified using ImageJ software for further analysis.

### Dual-Luciferase Reporter Assay

The binding site of PETPIR within the 3′ untranslated region (UTR) of YTHDF2 was predicted using the miRanda algorithm. Wild-type (WT) and mutant (MUT) 3′UTR sequences of YTHDF2 were cloned into the psiCHECK-2 plasmid vector (Promega, USA) to construct dual-luciferase reporter plasmids. HEK293T cells were seeded in 24-well plates at 1 × 10⁵ cells per well and cultured overnight in DMEM supplemented with 10% fetal bovine serum (FBS) at 37°C in a 5% CO₂ incubator. WT or MUT psiCHECK-2 plasmids (500 ng per well) were co-transfected with 50 nM PETPIR mimic or negative control (NC) using Lipofectamine 2000 (Invitrogen, Thermo Fisher Scientific) following the manufacturer’s protocol. Forty-eight hours post-transfection, luciferase activity was measured using the Dual-Luciferase Reporter Assay System (Promega, E1910). Firefly luciferase activity was normalized to Renilla luciferase activity (internal control). Relative luciferase activity, calculated as the ratio of firefly to Renilla luciferase signals, was detected using a BioTek microplate reader.

### m6A and mRNA Epitranscriptomic Microarray Analysis

The m6A epitranscriptomic profile of mRNA and lncRNA was analyzed using the Arraystar Human m6A-mRNA&lncRNA Epitranscriptomic Microarray (8×60K, Arraystar, Aksomics Inc.). Total RNA was extracted from the samples using TRIzol Reagent and quantified with a NanoDrop ND-1000 spectrophotometer (Thermo Scientific). RNA integrity was verified using an Agilent Bioanalyzer 2100 or MOPS denaturing electrophoresis to ensure high-quality samples. Total RNA underwent immunoprecipitation with an anti-N6-methyladenosine (m6A) antibody to isolate m6A-modified RNAs, which were captured on magnetic beads as the “IP” fraction. The unmodified RNAs in the supernatant were collected as the “Sup” fraction. RNAs from both fractions were labeled as complementary RNAs (cRNAs) with Cy5 (IP fraction) and Cy3 (Sup fraction) using Arraystar’s RNA labeling protocol. Labeled cRNAs were hybridized onto the microarray slides at 65°C for 17 hours. After hybridization, the slides were washed under stringent conditions and scanned using an Agilent G2505C Microarray Scanner to detect Cy5 and Cy3 fluorescence. Data acquisition was performed using Agilent Feature Extraction software (v11.0.1.1). Raw intensities for the IP (Cy5-labeled) and Sup (Cy3-labeled) fractions were normalized to the log2-scaled intensities of internal Spike-in RNAs. The “m6A quantity” for each RNA was calculated based on the normalized intensities of the IP fraction (Cy5-labeled). Differentially m6A-methylated RNAs between experimental groups were identified using predefined thresholds for fold change and statistical significance (p < 0.05).

### RNA Pull-Down Assay

RNA pull-down assays were performed using the RNA Pull-Down Kit (BersinBio, Guangzhou, China) according to the manufacturer’s instructions. Biotinylated NDRG1 RNA or its antisense RNA (negative control) was synthesized in vitro using the Biotin RNA Labeling Kit (Beyotime, China). The biotin-labeled RNA was incubated with whole-cell lysates from HTR-8/SVneo cells in RNA-binding buffer at 4°C for 4 hours with gentle rotation. RNA-protein complexes were captured using streptavidin-coated magnetic beads and washed extensively to remove nonspecific interactions. Bound proteins were eluted and analyzed by Western blot to identify RNA-interacting proteins. Specific antibodies were used to detect the target proteins in the pull-down samples.

### RNA Immunoprecipitation Assay

RNA immunoprecipitation (RIP) assays were conducted using the RIP Kit (BersinBio, Guangzhou, China) following the manufacturer’s protocol. Briefly, 2 × 10⁷ HTR-8/SVneo cells were lysed in RIP lysis buffer, and genomic DNA was removed to ensure RNA purity. The lysates were divided into three groups: input (reference control), immunoprecipitation (IP), and IgG control. The IP group lysates were incubated overnight at 4°C with specific antibodies against YTHDF2, while the IgG control group was treated with nonspecific IgG. Protein A/G magnetic beads were added to the samples and incubated at 4°C for 1 hour to isolate RNA-protein complexes. After extensive washing to remove unbound components, RNA was extracted from the complexes using TRIzol Reagent (Thermo Fisher Scientific). The purified RNA was reverse-transcribed into cDNA, and target RNA levels were quantified by qPCR. Data were normalized to the input control to correct for variations in sample handling and processing.

### Assessment of mRNA Stability

To assess mRNA stability, HTR-8/SVneo cells were treated with 5 µg/mL actinomycin D (MCE, HY-17559) to inhibit transcription. Cells were harvested at designated time points, and total RNA was extracted using TRIzol Reagent (Thermo Fisher Scientific) according to the manufacturer’s protocol. qRT-PCR was conducted to quantify the remaining levels of target mRNA at each time point, with normalization to an internal control. The relative mRNA abundance was expressed as a percentage of the baseline level (0-hour). The mRNA half-life was calculated using linear regression analysis of the logarithmic-transformed mRNA percentage over time.

#### Animal Experiments

All animal experiments were conducted in compliance with ethical guidelines and approved by the Animal Care and Use Committee of Nankai University.

Animal Model and Grouping: C57BL/6J female mice (8–12 weeks old) were mated with 10-week-old fertile males, and embryonic day 0.5 (E0.5) was defined as the day a vaginal plug was detected. Pregnant mice were randomly divided into three groups at E7.5 (n = 6 per group):

1) L-NAME group: Dams received tail vein injections of lentivirus carrying a negative control plasmid (LV-NC) (2 × 10⁷ infectious units/mouse, Hanbio, Shanghai, China) at E7.5, followed by daily subcutaneous injections of L-NAME (75 mg/kg/day, MedChemExpress, USA) from E7.5 to E17.5.

2) PETPIR group: Dams were injected with lentivirus carrying a PETPIR overexpression plasmid (LV-piR) (2 × 10⁷ infectious units/mouse) via the tail vein at E7.5 and received daily subcutaneous saline injections from E7.5 to E17.5.

3) NC group: Dams received lentivirus injections (LV-NC) and daily saline injections as described above.

Blood Pressure Measurement: Systolic blood pressure (SBP) was recorded daily from E0.5 to E18.5 using a non-invasive tail-cuff blood pressure system (Visitech Systems, BP2000). Mice were acclimated to the device for three days before measurements to minimize stress-induced variability.

Urine Collection and Protein Analysis: Urine samples were collected using metabolic cages over 24-hour periods at E6.5 and E17.5. Total protein and albumin concentrations were quantified with a bicinchoninic acid (BCA) protein assay kit (Beyotime Biotechnology, Shanghai, China). Daily protein excretion was calculated as the product of total protein concentration and 24-hour urine volume.

Tissue Collection and Analysis: On E18.5, pregnant dams were euthanized, and fetal and placental tissues were harvested. Placental weights were recorded, and tissues were either snap-frozen in liquid nitrogen for molecular studies or fixed in paraformaldehyde for histological analysis.

### Hematoxylin and Eosin Staining

Mice placental tissues were fixed in 4% paraformaldehyde at 4°C overnight, dehydrated through a graded ethanol series, and embedded in paraffin. Sections (4 μm thick) were prepared using a rotary microtome and stained with hematoxylin and eosin (H&E) for histological evaluation. Morphological changes in the placental architecture were analyzed under a light microscope, and high-resolution images were captured for further assessment.

### Quantitative Real-Time PCR and Western Blot Analysis

Total RNA was extracted from cells, tissues, and serum using TRIzol Reagent (Invitrogen, USA) according to the manufacturer’s instructions. cDNA synthesis was performed using the First Strand cDNA Synthesis Kit (Thermo Fisher Scientific, K1622). Quantitative real-time PCR (qRT-PCR) was conducted using Fast SYBR Green Master Mix (Applied Biosystems, Life Technologies, USA). U6 or ACTIN was used as the internal control for normalization. Relative gene expression levels were calculated using the 2-ΔΔCt method.

Total protein was extracted from cells or tissues, and the protein concentration was quantified using a BCA Protein Assay Kit (Beyotime Biotechnology, Shanghai, China). Equal amounts of protein were separated by SDS-PAGE and transferred onto Immobilon-PSQ PVDF membranes (Millipore). Membranes were blocked with 5% non-fat milk in TBST for 2 hours at room temperature, followed by overnight incubation at 4°C with specific primary antibodies. After three washes with TBST, membranes were incubated with HRP-conjugated secondary antibodies for 1 hour at room temperature. Protein bands were visualized using an enhanced chemiluminescence (ECL) detection system.

### Statistical analysis

Statistical analyses were performed using GraphPad Prism software. Comparisons between the two groups were conducted using an unpaired t-test. For multiple group comparisons, one-way or two-way analysis of variance (ANOVA) was applied, followed by Bonferroni’s post-hoc test to assess specific group differences. Data are presented as mean ± standard error (SE). A p-value < 0.05 was considered statistically significant.

## Results

### Clinical Characteristics and Identification of Preeclampsia-Associated piRNAs

A total of 29 pregnant women diagnosed with preeclampsia (PE group) and 29 healthy controls (Ctl group) were included in this study. The clinical characteristics of the participants are summarized in Table 1. Maternal age and body mass index (BMI) were comparable between the two groups, with no statistically significant differences (maternal age: 31.3 ± 3.1 years in the PE group vs. 32.1 ± 4.6 years in the Ctl group; BMI: 28.9 ± 6.1 kg/m² in the PE group vs. 26.7 ± 4.2 kg/m² in the Ctl group). The mean gestational age at delivery also showed no significant difference (246.4 ± 21.1 days for the PE group vs. 248.8 ± 19.7 days for the Ctl group).

In contrast, hallmark clinical features of preeclampsia were observed in the PE group, including significantly elevated blood pressure and proteinuria. The systolic (162.9 ± 17.4 mmHg) and diastolic (101.1 ± 11.5 mmHg) blood pressures in the PE group were markedly higher than those in the Ctl group (110.8 ± 7.7 mmHg and 67.9 ± 5.6 mmHg, respectively; p < 0.05). Proteinuria was evident in the PE group, with an average 24-hour urinary protein level of 2.8 ± 2.0 g, while no proteinuria was detected in the Ctl group (p < 0.05). Furthermore, adverse fetal outcomes were noted in the PE group, with significantly lower infant birth weights (2087 ± 663 g vs. 2553 ± 734 g in the Ctl group; p < 0.05).

To investigate the role of piRNAs in preeclampsia, placental samples from four preeclamptic (PE) and four control (Ctl) subjects were analyzed using piRNA microarray profiling (Figure 1A). A total of 935 piRNAs were significantly upregulated, and 1538 piRNAs were significantly downregulated in the PE group compared to the Ctl group (Figure 1B). Among these, DQ601040 emerged as the most significantly upregulated piRNA (Figure 1C). Validation in larger samples confirmed the differential expression patterns of key piRNAs, with consistent upregulation of DQ601040 in preeclamptic placentas (Figures 1D–G). Further qRT-PCR analysis of serum samples revealed significantly elevated DQ601040 levels in preeclamptic subjects compared to controls (Figure 1H). Receiver operating characteristic (ROC) curve analysis demonstrated a strong diagnostic potential for serum DQ601040 levels, yielding an area under the curve (AUC) of 0.9917 (95% CI: 0.9654–1.000) (Figure 1I). Additionally, cross-species conservation analysis showed that DQ601040 is highly conserved across humans and other mammals (Figure 1J), suggesting its potential functional importance in preeclampsia pathophysiology.

**Figure 1.**
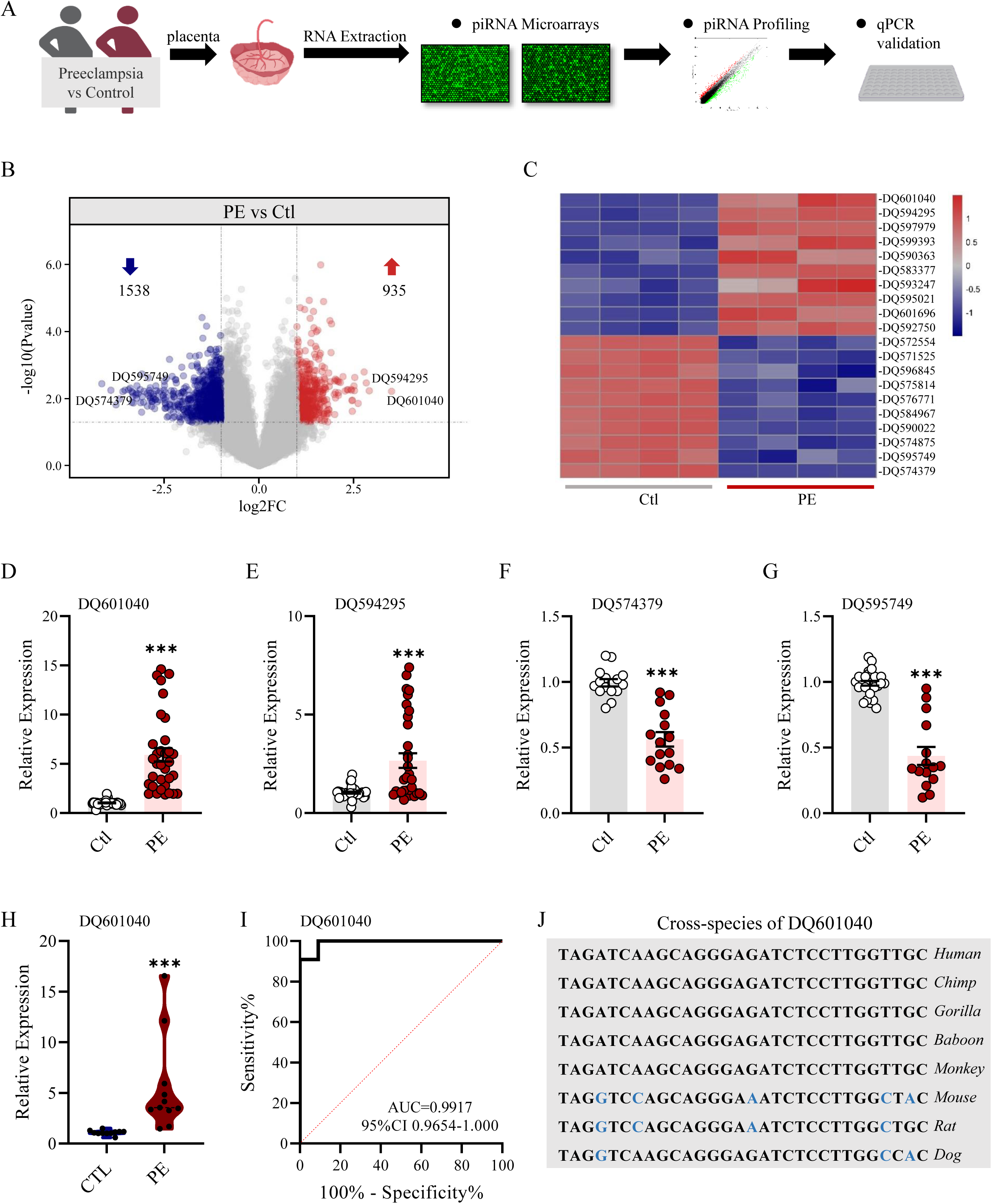
Identification and characterization of PETPIR in preeclamptic placentas. (A) Schematic representation of the experimental workflow used to identify differentially expressed piRNAs in preeclamptic (PE) versus control (Ctl) placentas. Placental samples were collected, RNA was extracted, piRNA profiles were analyzed via microarray, and selected candidates were validated using qRT-PCR. (B) Volcano plot displaying differentially expressed piRNAs between PE and Ctl placentas. Red dots indicate upregulated piRNAs (476), and blue dots represent downregulated piRNAs (839). (C) Heatmap of the top 20 differentially expressed piRNAs in PE versus Ctl placentas. Each column represents an individual sample, while rows show the expression levels of specific piRNAs. Red and blue indicate higher and lower expression levels, respectively. (D-G) qRT-PCR validation of selected piRNAs, including DQ601040, DQ594295, DQ574379, and DQ597849, in larger samples. (H) Violin plot demonstrating the relative expression of PETPIR in the blood serum of PE and Ctl groups. (I) Receiver operating characteristic (ROC) curve for DQ601040 showing its diagnostic potential in distinguishing PE from Ctl blood serum. (J) Cross-species sequence alignment of DQ601040 highlighting its evolutionary conservation across multiple species, including humans, primates, and rodents. Data are presented as mean ± SEM. ***p < 0.001 vs control (Ctl) group.

### PETPIR Suppresses Trophoblast Cell Proliferation, Migration, and Invasion

To investigate the cellular localization of PETPIR, we performed RNA fluorescence in situ hybridization (FISH) in HTR-8/SVneo trophoblast cells. The analysis revealed that PETPIR is distributed in both the nucleus and cytoplasm (Figure 2A, B). Given its high expression in trophoblast cells and its potential functional relevance, we designated DQ601040 as Preeclampsia-Associated Trophoblast piRNA (PETPIR).

**Figure 2.**
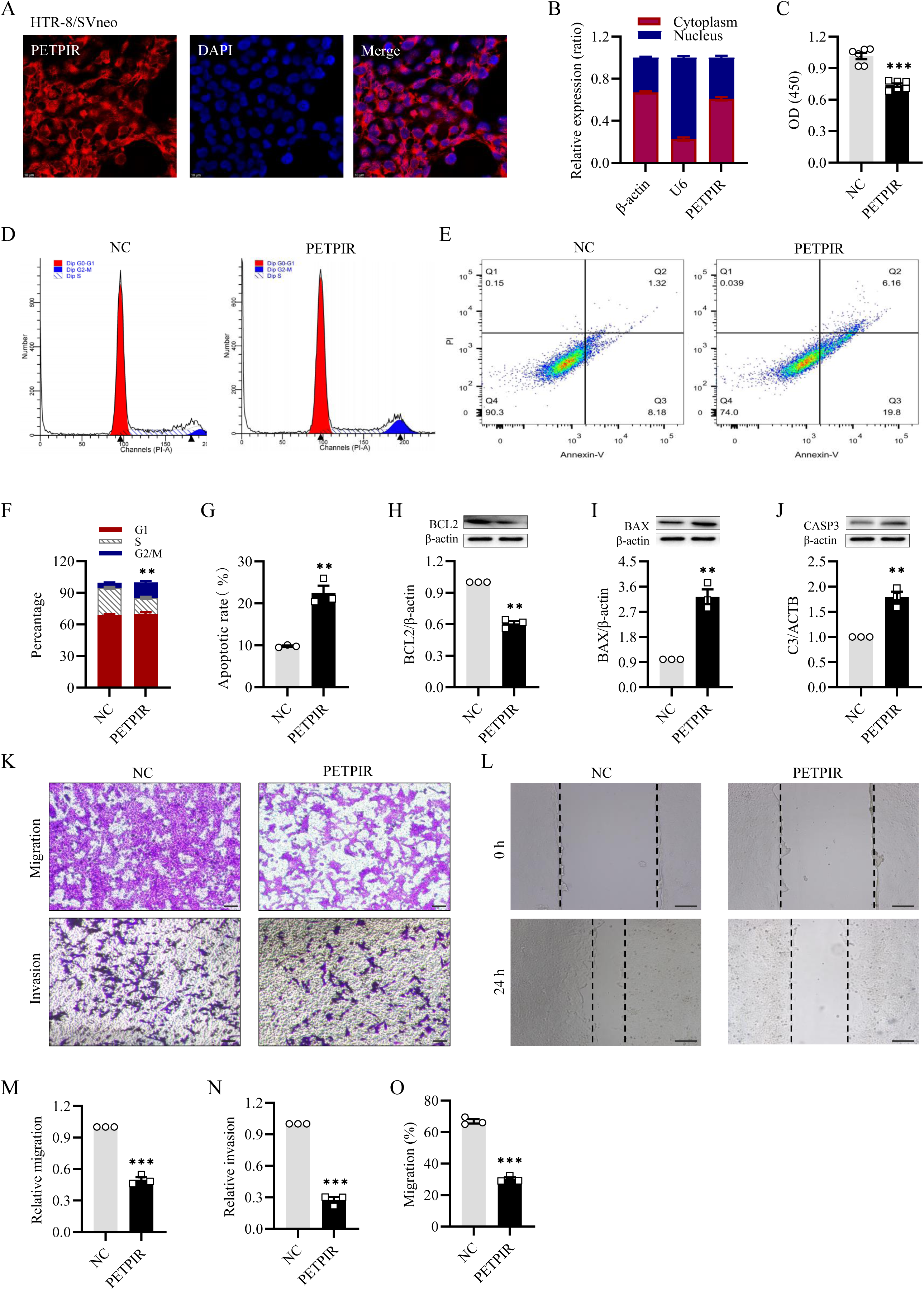
PETPIR overexpression suppresses trophoblast cell proliferation, induces apoptosis, and inhibits migration and invasion. (A) Immunofluorescence staining showing PETPIR localization in HTR8/SVneo cells. Scale bar, 10μm. (B) Quantitative analysis of PETPIR subcellular localization, indicating significant enrichment in the cytoplasm compared to the nucleus. (C) CCK-8 assay demonstrating reduced cell viability in PETPIR-overexpressing cells compared to the negative control (NC) group. (D) Flow cytometry analysis of the cell cycle distribution shows that PETPIR overexpression induces G0/G1 phase arrest, with a corresponding decrease in the S-phase population. (E) Apoptosis analysis by Annexin V/PI staining revealed a significant increase in both early and late apoptotic cell populations in PETPIR-overexpressing cells compared to the NC group. (F) Quantitative data confirmed a higher percentage of cells in the G0/G1 phase and reduced S-phase entry in PETPIR-overexpressing cells. (G-J) Western blot analysis of apoptosis-related proteins. PETPIR overexpression increases the levels of pro-apoptotic markers Bax (I) and Casp3 (J) while decreasing the expression of the anti-apoptotic protein Bcl-2 (H). (K) Representative images of transwell migration and invasion assays show that PETPIR overexpression significantly reduces the number of migrating and invading trophoblast cells compared to the NC group. Scale bar, 50μm. (L) Wound healing assay demonstrating impaired wound closure in PETPIR-overexpressing cells at 24 hours compared to the NC group. Scale bar, 200μm. (M-O) Quantitative analyses of transwell migration (M), invasion (N), and wound healing (O), confirm significant reductions in motility and invasive capacity in PETPIR-overexpressing cells. Data are presented as mean ± SEM. **p < 0.01, ***p < 0.001 vs Negative control (NC) group.

To explore the functional role of PETPIR in trophoblast cells, we overexpressed PETPIR and assessed its effects on cell proliferation, cell cycle progression, and apoptosis. The CCK-8 assay revealed a significant reduction in cell viability following PETPIR overexpression compared to the negative control (NC) group (p < 0.001, Figure 2C). Cell cycle analysis using flow cytometry demonstrated that PETPIR overexpression induced G1 phase arrest, with a significant increase in the proportion of cells in the G1 phase and a corresponding decrease in the S phase (p < 0.01, Figure 2D, F). Apoptosis analysis showed elevated apoptotic rates in PETPIR-overexpressing cells compared to the NC group (p < 0.01, Figure 2E, G). These findings were supported by Western blot results, which showed increased expression of pro-apoptotic markers, including BAX, and cleaved caspase-3, and a reduction in the anti-apoptotic marker BCL2 in the PETPIR group (p < 0.01, Figure 2H–J).

To further examine the impact of PETPIR on trophoblast migration and invasion, we performed Transwell and wound healing assays. Transwell assays revealed a significant decrease in both migration and invasion capabilities in PETPIR-overexpressing cells (p < 0.001, Figure 2K, M, N). Similarly, the wound healing assay demonstrated impaired wound closure in the PETPIR group compared to the NC group (p < 0.001, Figure 2L, O). These results indicate that PETPIR plays a critical inhibitory role in trophoblast cell proliferation, migration, and invasion while promoting apoptosis and cell cycle arrest.

### PETPIR Activates the Hypoxia Signaling Pathway in Trophoblast Cells

To elucidate the molecular mechanisms underlying PETPIR’s effects on trophoblast cells, RNA sequencing (RNA-seq) was conducted on HTR-8/SVneo cells overexpressing PETPIR. Differential gene expression analysis identified 396 upregulated and 111 downregulated genes in PETPIR-overexpressing cells compared to the NC group (Figures 3A, B).

**Figure 3.**
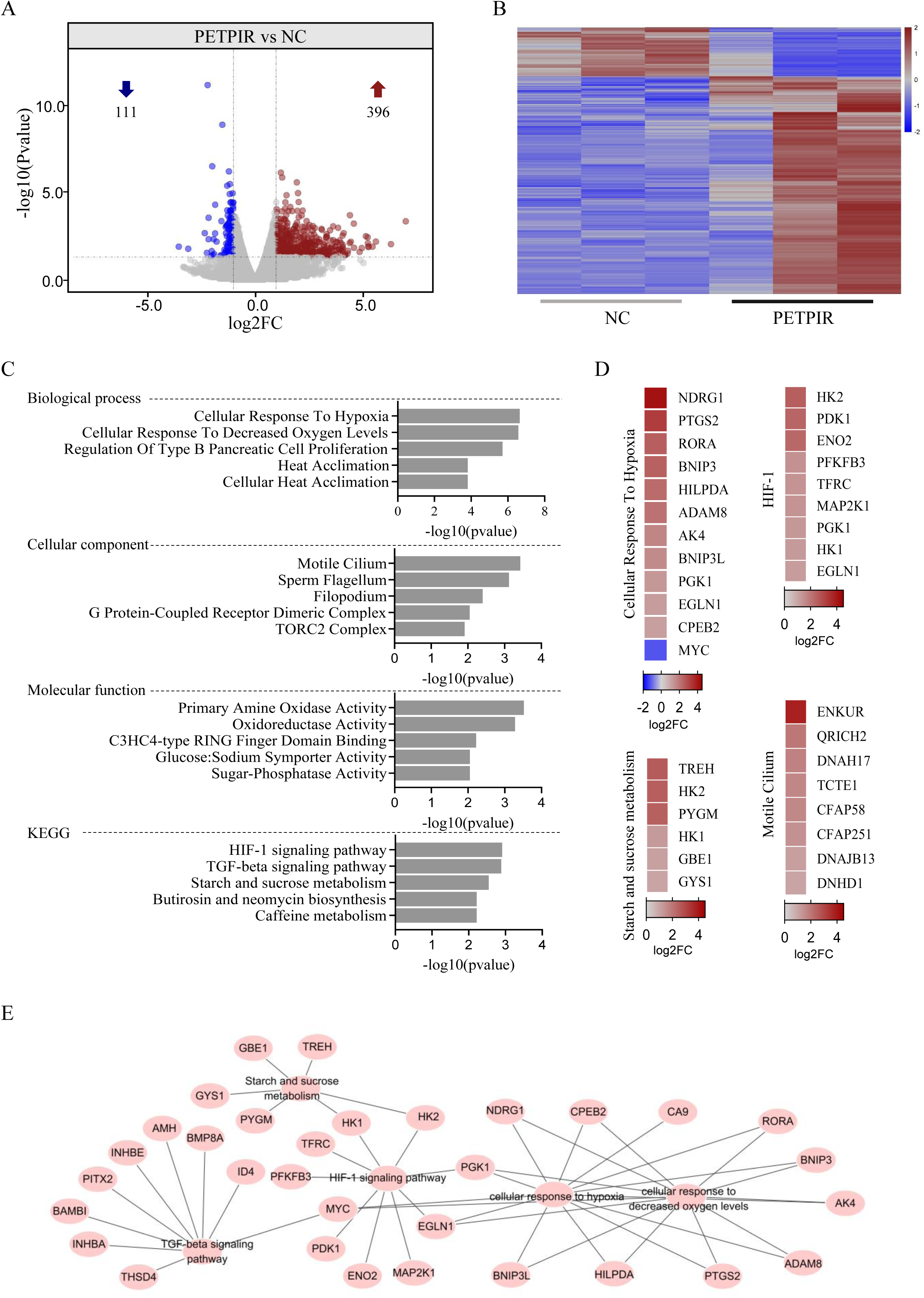
PETPIR overexpression alters transcriptomic profiles and activates hypoxia-responsive pathways in trophoblast cells. (A) Volcano plot illustrating differentially expressed genes (DEGs) in PETPIR-overexpressing HTR8/SVneo cells compared to the negative control (NC) group. A total of 396 genes are upregulated (red), and 111 genes are downregulated (blue). (B) Heatmap depicting the expression profiles of DEGs in PETPIR-overexpressing cells, showing distinct transcriptomic alterations compared to the NC group. Each column represents a sample, and each row represents a DEG, with red and blue indicating upregulation and downregulation, respectively. (C) Gene Ontology (GO) enrichment analysis of DEGs. (D) Heatmap showing the expression levels of key genes associated with hypoxia response, HIF-1, Starch and sucrose metabolism, and Motile Cilium. (E) Gene interaction network analysis focused on the HIF-1 signaling pathway and related hypoxia-responsive genes.

Gene Ontology (GO) enrichment analysis of these differentially expressed genes (DEGs) revealed significant enrichment in biological processes related to cellular responses to hypoxia and decreased oxygen levels. Cellular components such as the motile cilium and TORC2 complex were prominently represented, while molecular function analysis showed enrichment in primary amine oxidase activity and oxidoreductase activity (Figure 3C). Kyoto Encyclopedia of Genes and Genomes (KEGG) pathway analysis further identified the hypoxia-inducible factor-1 (HIF-1) signaling pathway as the most significantly enriched, alongside the TGF-beta signaling pathway and oxidative stress-related metabolic pathways (Figure 3C). These findings suggest that PETPIR regulates critical hypoxia-driven processes in trophoblast cells. A gene interaction network analysis further highlighted key nodes associated with hypoxia signaling, including NDRG1, PGK1, EGLN1, CPEB2, HK1, and PFKFB3, which are important mediators of cellular responses to hypoxia (Figure 3D). These results demonstrate that PETPIR activates the hypoxia-associated signaling pathway and its critical role in regulating trophoblast cell responses to hypoxia.

### PETPIR Modulates m6A Methylation by Targeting YTHDF2

Emerging evidence highlights the regulatory role of piRNAs in m6A RNA methylation through interactions with m6A writers, erasers, and readers. To determine whether PETPIR influences m6A modifications, dot blot assays were performed, revealing a significant increase in global m6A levels in the PETPIR-overexpressing HTR8/SVneo cells compared to the NC group (Figures 4A, B). These findings indicate that PETPIR is actively involved in m6A-mediated regulatory pathways.

**Figure 4.**
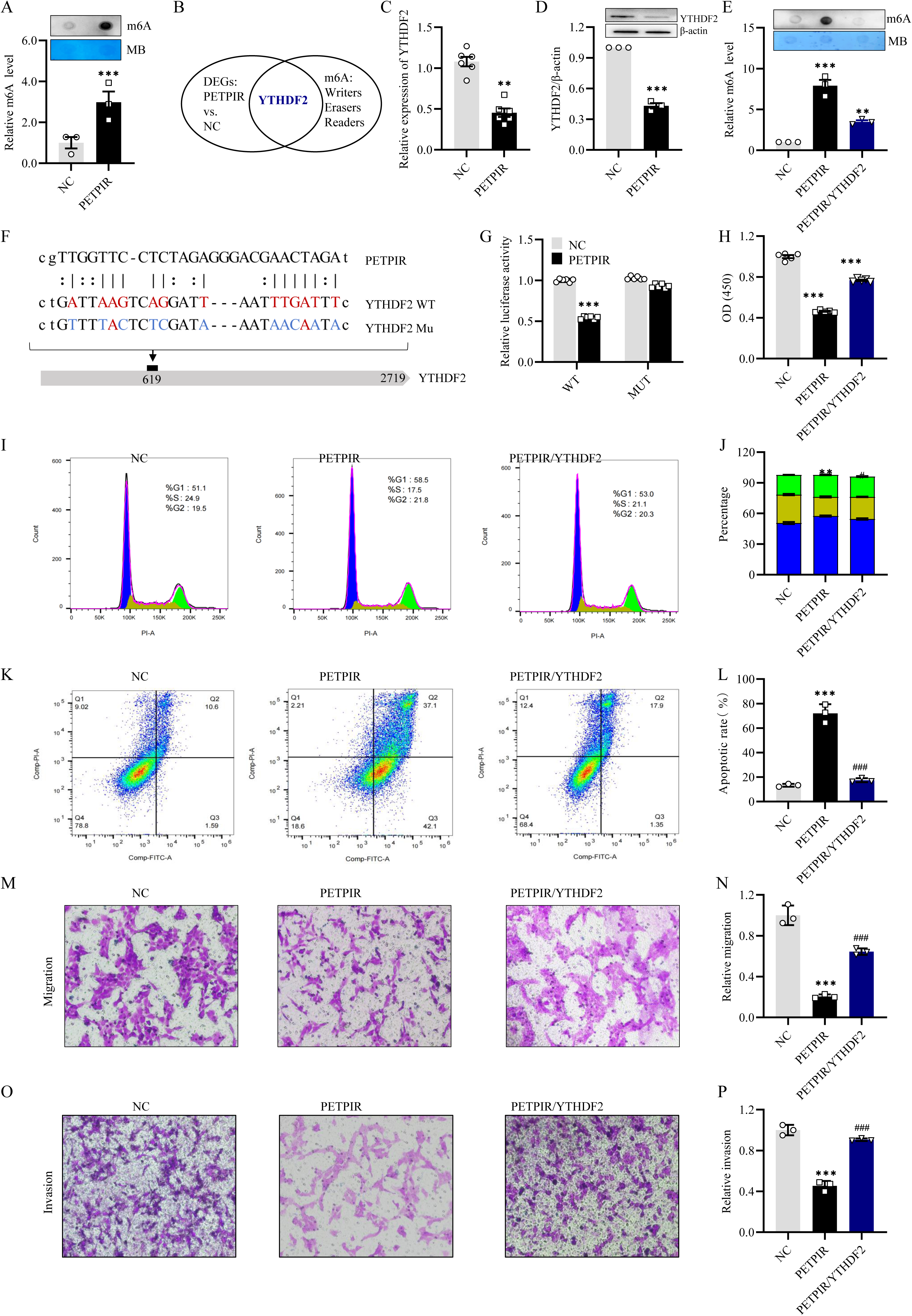
PETPIR enhances m6A methylation through its interaction with YTHDF2. (A) Dot blot analysis of global m6A methylation levels in HTR8/SVneo cells. PETPIR-overexpressing cells exhibit significantly higher m6A levels compared to the negative control (NC) group. (B) Venn diagram depicting the overlap of differentially expressed genes (DEGs) in PETPIR-overexpressing cells and the writers, readers, and erasers of m6A. (C, D) RT-PCR and Western blot analysis demonstrating upregulation of YTHDF2 at the mRNA (C) and protein (D) levels in PETPIR-overexpressing cells. (E) Dot blot assay showing m6A methylation levels in PETPIR-overexpressing cells with or without YTHDF2 co-expression. YTHDF2 is essential for the PETPIR-induced increase in m6A methylation. (F) Schematic representation of YTHDF2 functional domains, highlighting the RNA-binding region responsible for PETPIR interaction. Mutations at key m6A consensus motifs (red) disrupt this interaction. (G) Relative luciferase activity of wild-type (WT) or mutant (Mut) YTHDF2 constructs in the presence of PETPIR. PETPIR specifically interacts with WT YTHDF2, but not with the mutant, indicating a sequence-specific interaction. (H) CCK-8 assay assessing cell viability in response to PETPIR overexpression and YTHDF2 co-expression. (I) Cell cycle analysis by flow cytometry. PETPIR overexpression induces G0/G1 phase arrest in HTR8/SVneo cells, leading to a reduction in the S phase population. (J) Quantification of cell cycle distribution across G0/G1, S, and G2/M phases. YTHDF2 significantly reverses the PETPIR-induced G0/G1 arrest. (K) Flow cytometry analysis of apoptosis using Annexin V/PI staining. PETPIR overexpression markedly increases the percentage of apoptotic cells, an effect partially rescued by YTHDF2 co-expression. (L) Quantification of apoptotic cell populations from panel (K). (M) Representative images of cell migration assessed by transwell migration assays. (N) Quantification of migrated cells from panel (M). Scale bar, 50μm. (O) Representative images of cell invasion assessed by transwell invasion assays. Scale bar, 50μm. (P) Quantification of invaded cells from panel (O). Data are presented as mean ± SEM. *p < 0.05, **p < 0.01, ***p < 0.001 vs Negative control (NC) group. ###p < 0.001 vs PETPIR group.

To investigate the underlying mechanism, RNA-seq analysis of differentially expressed genes in PETPIR-overexpressing cells identified the m6A reader protein YTHDF2 as significantly downregulated (Figure 4C). RT-PCR and Western blot analyses confirmed a marked decrease in YTHDF2 expression at both the mRNA and protein levels in PETPIR-overexpressing cells (Figures 4D, E). To determine whether PETPIR directly interacts with YTHDF2, bioinformatic analysis using the miRanda algorithm predicted a direct binding site for PETPIR within the YTHDF2 mRNA (Figure 4G). This interaction was validated using a dual-luciferase reporter assay. Cells co-transfected with the PETPIR mimic and YTHDF2 wild-type (WT) reporter plasmid exhibited a significant reduction in luciferase activity, while mutation of the predicted seed sequence in the YTHDF2 mutant (MU) reporter abolished this effect, confirming the specificity of PETPIR binding to YTHDF2.

Further functional analysis demonstrated that YTHDF2 plays a crucial role in PETPIR-mediated m6A regulation. Overexpression of YTHDF2 in PETPIR-overexpressing cells significantly attenuated the increase in global m6A levels induced by PETPIR (Figures 4H, I), suggesting the key role of YTHDF2 as a mediator of PETPIR’s effects on m6A methylation. To evaluate whether YTHDF2 mediates the effects of PETPIR on trophoblast cell function, HTR8/SVneo cells were co-transfected with PETPIR overexpression constructs and YTHDF2 overexpression plasmids. Functional assays revealed that PETPIR overexpression significantly reduced trophoblast cell proliferation, an effect that was reversed by YTHDF2 overexpression (Figure 4J). Flow cytometry analysis showed increased apoptosis in PETPIR-overexpressing cells, which was mitigated by YTHDF2 overexpression (Figure 4K-4L). Additionally, Transwell migration and invasion assays demonstrated reduced migratory and invasive capacities in PETPIR-overexpressing cells, both of which were restored by YTHDF2 co-expression (Figure 4M-4P). These results establish the critical role of YTHDF2 in mediating the functional effects of PETPIR in trophoblast cells, highlighting the importance of the PETPIR-YTHDF2 axis in regulating m6A methylation and trophoblast cell function.

### Transcriptome-Wide Analysis of RNA m6A Methylation in Preeclampsia

Given our finding that PETPIR modulates m6A methylation in trophoblast cells, we extended our investigation to a transcriptome-wide analysis of RNA m6A methylation in placental tissues from women with preeclampsia (PE) and those with normal pregnancies (Ctl) (Figure 5A). This analysis aimed to identify global epitranscriptomic changes associated with PE. The results revealed significant alterations in m6A methylation patterns and gene expression between the two groups. Specifically, 444 upregulated and 605 downregulated genes were identified in preeclamptic placentas (Figure 5B). Gene Ontology (GO) analysis of differentially expressed genes (DEGs) in the PE group showed significant enrichment in processes such as “tube closure,” “hormone receptor binding,” and “growth factor activity” (Figure 5D, 5E).

**Figure 5.**
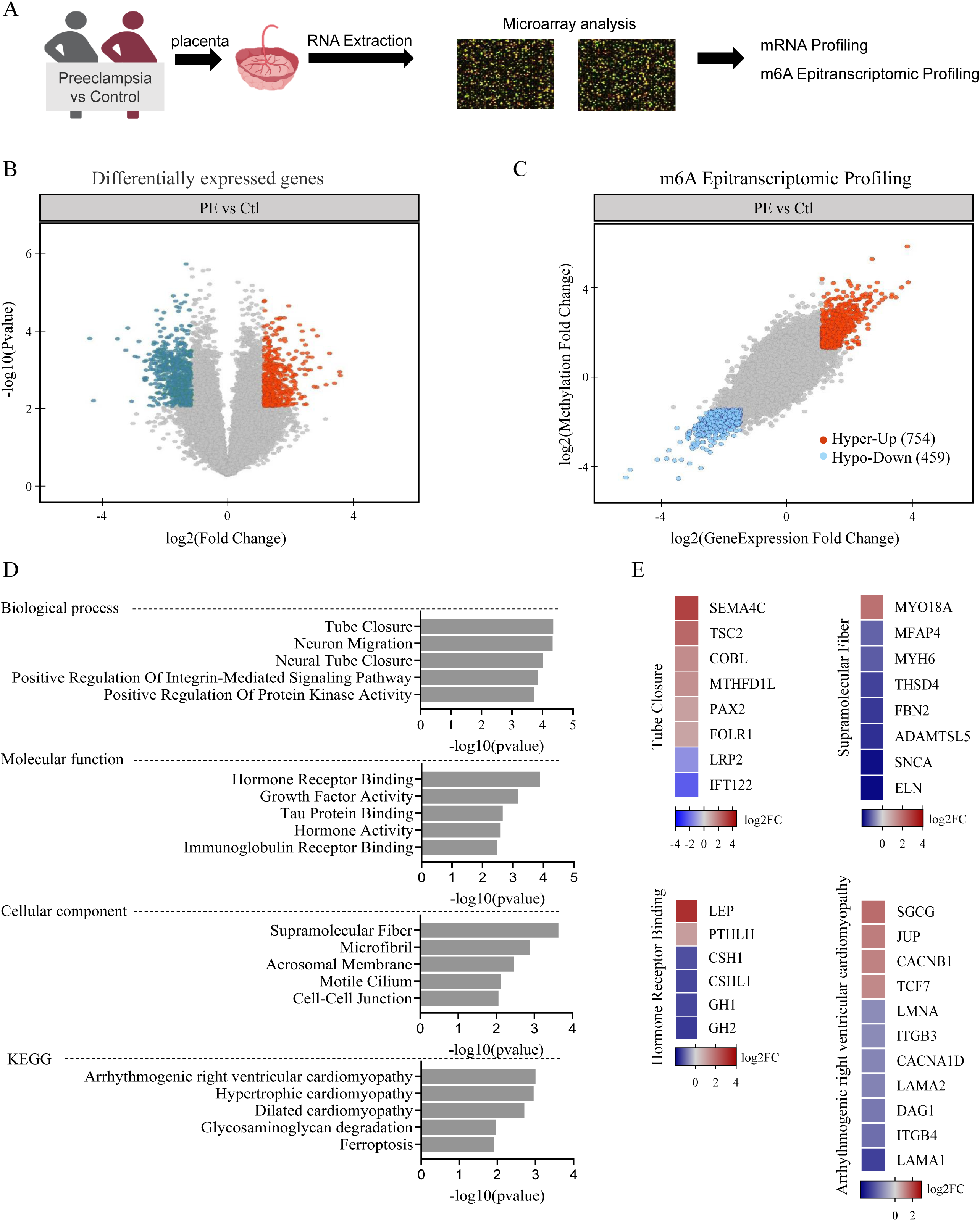
Transcriptome-wide m6A epitranscriptomic analysis reveals differential methylation in preeclampsia. (A) Schematic representation of the experimental workflow for mRNA and m6A epitranscriptomic profiling in placental samples from preeclampsia (PE) and control (Ctl) groups. (B) Volcano plot illustrating differentially expressed genes (DEGs) between PE and Ctl groups. Upregulated genes are shown in red, and downregulated genes are in blue. (C) m6A epitranscriptomic profiling identifies 754 hypermethylated (Hyper-Up) and 459 hypomethylated (Hypo-Down) genes in PE placentas compared to controls, indicating significant epitranscriptomic alterations. (D) Gene Ontology (GO) enrichment analysis of DEGs. (E) Heatmaps showing key genes enriched in critical biological pathways.

A total of 754 hypermethylated (Hyper-Up) and 459 hypomethylated (Hypo-Down) genes were identified in PE placentas compared to controls (Figure 5C). Enrichment analysis of hypermethylated genes revealed their involvement in key biological processes, including “tube closure,” “tropomyosin binding,” and “apical junction complexes,” which are critical for cell communication and trophoblast function (Figure 6A). Disease association analysis (Jensen diseases) linked hypermethylated genes to “hypertension” and “cerebrovascular disease,” emphasizing their potential role in the pathogenesis of PE. A gene interaction network for hypermethylated genes identified key regulatory hubs, including IL6 and VEGFA, highlighting their central roles in hypoxia, angiogenesis, and inflammation (Figures 6B, C).

**Figure 6.**
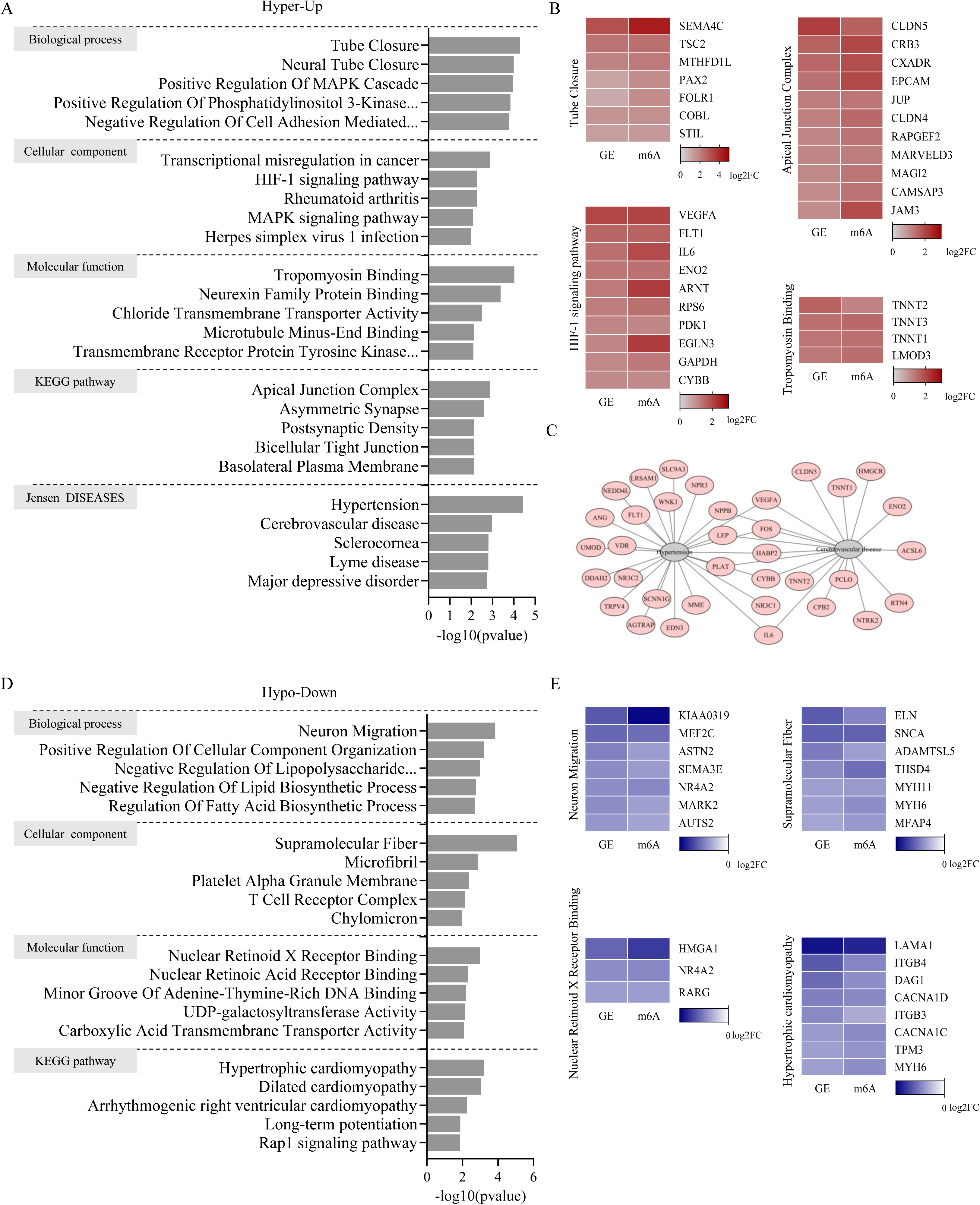
Functional and pathway enrichment analysis of m6A-modified genes in preeclampsia. (A) Gene Ontology (GO) enrichment analysis of hypermethylated and upregulated genes (Hyper-Up) in preeclamptic placentas. (B) Heatmaps of key genes involved in tube closure, apical junction complexes, and HIF-1 signaling, demonstrating significant differential expression and methylation in Hyper-Up genes in preeclamptic placentas. (C) Gene interaction network of hypermethylated genes in hypertension and cardiovascular disease. (D) GO enrichment analysis of hypomethylated and downregulated genes (Hypo-Down). (E) Heatmaps of critical genes involved in neuron migration, supramolecular fiber, nuclear retinoid X receptor binding, and hypertrophic cardiomyopathy.

In contrast, hypomethylated genes (Hypo-Down) were enriched in biological processes such as “neuron migration,” “positive regulation of cell component organization,” and “regulation of lipid biosynthetic processes.” Cellular component analysis revealed significant associations with “supramolecular fibers” and “platelet alpha granule membranes,” while molecular function analysis indicated enrichment in “nuclear retinoid receptor binding” and “carboxylic acid transmembrane transporter activity” (Figure 6D). Pathway analysis of hypomethylated genes highlighted significant associations with “cardiomyopathy,” “Ras signaling,” and “long-term potentiation,” suggesting disruptions in metabolic and signaling pathways essential for normal placental development. Genes such as HMGA1, ELN, and LAMA1 were prominent among the hypomethylated and downregulated genes (Figure 6E). These findings reveal substantial epitranscriptomic alterations in RNA m6A methylation in PE, with hypermethylation predominantly driving hypoxia- and inflammation-related pathways, and hypomethylation contributing to metabolic and structural dysregulation.

### NDRG1 is a Key Downstream Target of the PETPIR/YTHDF2 Axis

To identify potential downstream targets of the PETPIR/YTHDF2 axis, we performed an integrative analysis by overlapping the 754 hypermethylated and upregulated genes identified in preeclampsia (PE) placentas with the 355 upregulated genes from RNA-seq analysis of PETPIR-overexpressing HTR8/SVneo cells. This analysis identified 20 shared genes, with NDRG1 emerging as the most significantly upregulated (Figures 7A, B).

**Figure 7.**
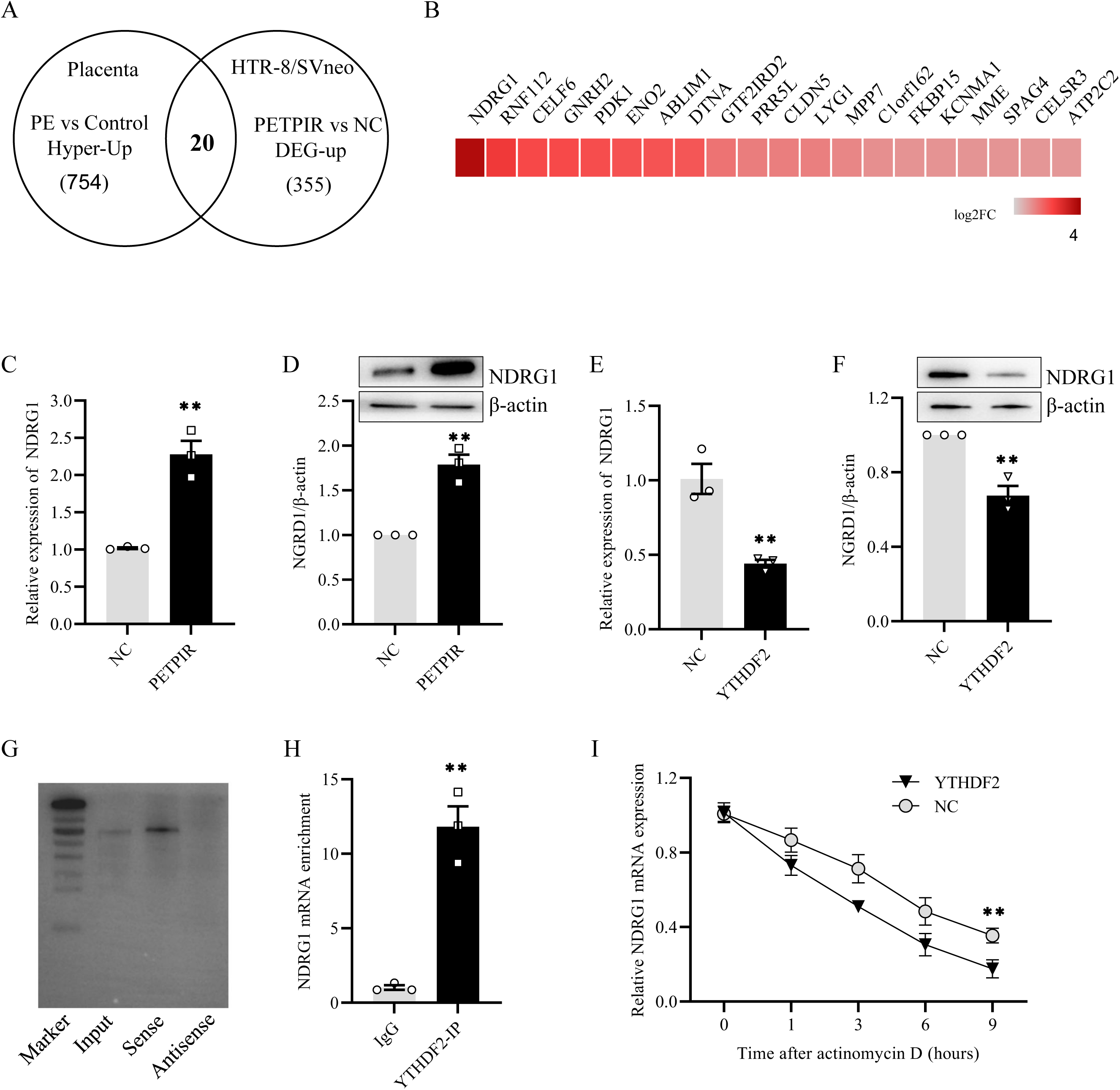
NDRG1 is identified as a downstream target of the PEPIR/YTHDF2 axis. (A) Venn diagram illustrating the overlap of hypermethylated and upregulated genes in preeclamptic placentas and upregulated differentially expressed genes (DEGs) in PEPIR-overexpressing HTR8/SVneo cells. Twenty shared genes were identified, including NDRG1 as the top candidate. (B) Heatmap depicting the expression levels of the 20 shared genes, highlighting NDRG1 as the most significantly upregulated gene. (C, D) Quantitative PCR (qPCR) and Western blot analyses confirmed significant upregulation of NDRG1 at both mRNA and protein levels in PEPIR-overexpressing cells compared to negative control (NC) cells. (E, F) Overexpression of YTHDF2 leads to a significant reduction in NDRG1 expression at both mRNA and protein levels. (H) RNA pull-down assay confirming the direct interaction between NDRG1 mRNA and YTHDF2, with significant enrichment in the antisense control lane. (I) RNA immunoprecipitation (RIP) assay showing significant enrichment of NDRG1 transcripts in YTHDF2 immunoprecipitates compared to IgG controls, validating the interaction. (J) Actinomycin D chase assay demonstrates that YTHDF2 overexpression decreases NDRG1 mRNA stability by slowing its degradation rate compared to NC cells. Data are presented as mean ± SEM. **p < 0.01 vs Negative control (NC) group.

Quantitative PCR and Western blot analyses validated that NDRG1 expression was significantly increased at both the mRNA and protein levels in PETPIR-overexpressing cells compared to the NC group (Figures 7C, D). Overexpression of YTHDF2 resulted in reduced NDRG1 expression at both the mRNA and protein levels, demonstrating the regulatory link between YTHDF2 and NDRG1 (Figures 7E, F). To investigate whether YTHDF2 directly regulates NDRG1 mRNA, RNA immunoprecipitation (RIP) assays using an anti-YTHDF2 antibody were performed. The results showed significant enrichment of NDRG1 transcripts in YTHDF2-immunoprecipitated fractions compared to IgG controls, confirming that YTHDF2 directly binds to NDRG1 mRNA. RNA pull-down assays further confirmed this interaction, showing enriched binding of YTHDF2 to NDRG1 mRNA in the YTHDF2-bound RNA fraction compared to controls (Figure 7G). RIP-qPCR results also demonstrated significant enrichment of NDRG1 transcripts in YTHDF2 immunoprecipitates, reinforcing the hypothesis of direct interaction (Figure 7H). To explore the functional impact of YTHDF2-mediated regulation on NDRG1 mRNA stability, actinomycin D chase assays were conducted. YTHDF2 overexpression significantly decreased the stability of NDRG1 mRNA (Figure 7I), indicating that YTHDF2-mediated m6A methylation stabilizes NDRG1 transcripts. These results identify NDRG1 as a critical downstream target of the PETPIR/YTHDF2 axis.

### Overexpression of PETPIR Induces Preeclampsia-Like Syndrome in Mice

To investigate the in vivo role of PETPIR, we established a mouse model of preeclampsia-like syndrome by overexpressing PETPIR in pregnant mice. L-NAME treatment, a well-established method for inducing preeclampsia, was used as a positive control (Figure 8A). Quantitative RT-PCR analysis confirmed a significant increase in PETPIR expression in the placental tissues of the PETPIR-overexpressing group, compared to the control group (Figure 8B).

**Figure 8.**
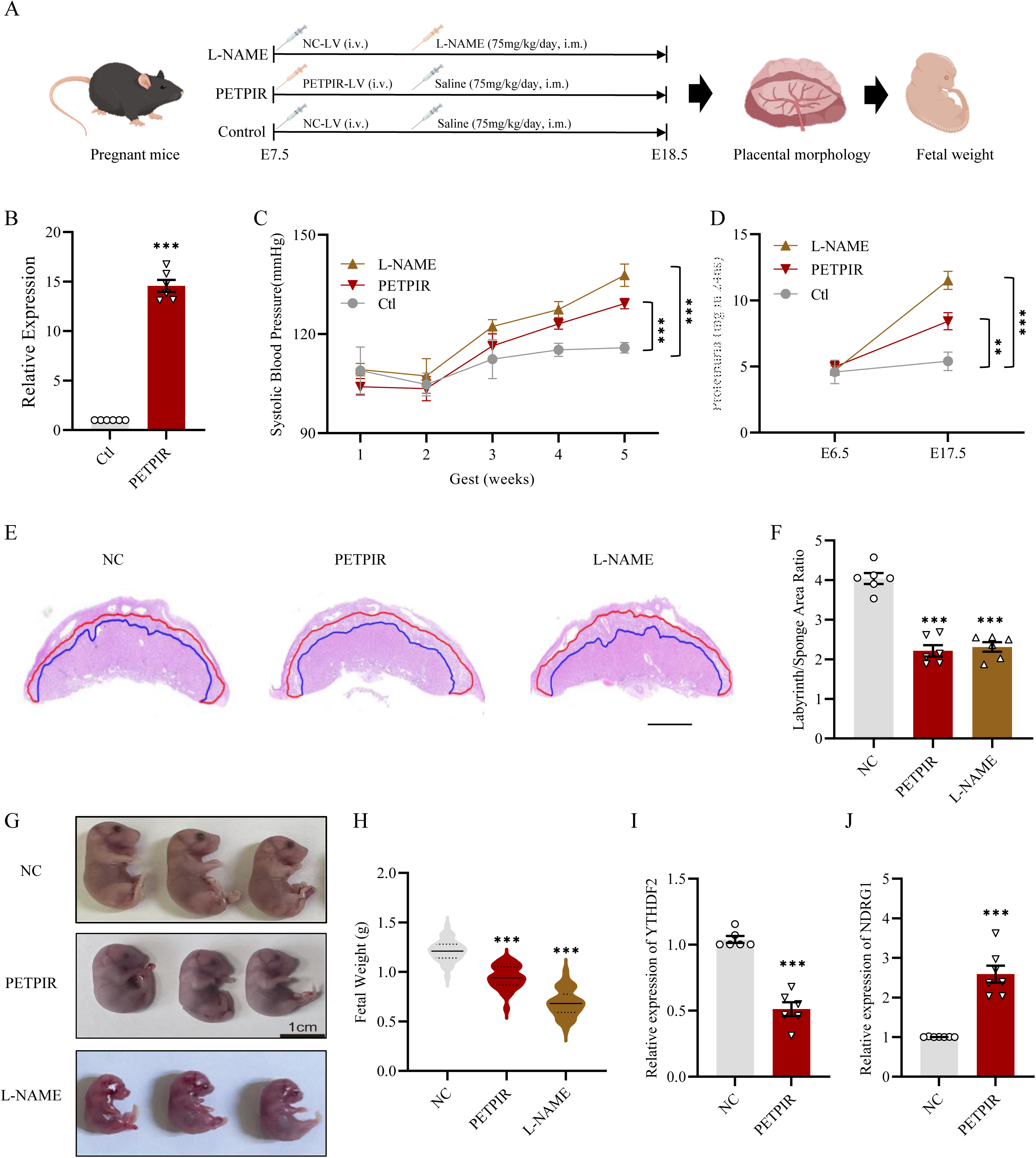
PEPIR overexpression induces a preeclampsia-like phenotype in a mouse model. (A) Schematic representation of the experimental design. Pregnant mice were injected with either PEPIR-overexpressing plasmids, L-NAME (a positive control), or saline (negative control, NC). Placental morphology and fetal outcomes were analyzed on embryonic day 18.5 (E18.5). (B) Quantitative RT-PCR analysis demonstrating significant upregulation of PEPIR in placentas from PEPIR-overexpressing mice compared to the NC group. (C) Systolic blood pressure (SBP) measurements showed significantly elevated SBP in PEPIR-overexpressing mice, similar to the L-NAME-treated group, during gestation. (D) Measurement of Proteinuria at E6.5 and E17.5 in three groups. (E) Histological images of placentas (H&E staining) from NC, L-NAME-treated, and PEPIR-overexpressing groups. Scale bar = 100 μm. (F) Quantification of lumen-to-wall ratios indicates significant reductions in the PEPIR and L-NAME groups. (G) Gross morphology of fetuses demonstrating smaller reduced fetal weights (H) in the PEPIR-overexpressing and L-NAME-treated groups compared to NC mice. RT-PCR analysis shows significant downregulation of YTHDF2 and (I) upregulation of NDRG1 (J) in placentas from PEPIR-overexpressing mice, implicating its role in the observed pathological changes associated with preeclampsia.

Systolic blood pressure measurements during gestation revealed a marked elevation in the PETPIR group, comparable to the levels observed in the L-NAME-treated group, and significantly higher than in the NC group (Figure 8C). Additionally, the PETPIR and L-NAME-treated groups exhibited elevated protein levels in the urine, a hallmark of preeclampsia (Figure 8D). Histological analysis of uteroplacental sections showed a reduced area ratio of the labyrinth layer to the sponge layer in the PETPIR-overexpressing group, resembling the pathological changes observed in the L-NAME-treated group (Figure 8E). The lumen-to-wall ratio of spiral arteries was significantly reduced in the PETPIR group (Figure 8F). At E18.5, placental morphology and fetal weights were assessed. PETPIR mice displayed significantly reduced fetal weights compared to the NC group (Figures 8G, H). Furthermore, RT-PCR analysis revealed significant downregulation of YTHDF2 (Figure 8I) and upregulation of NDRG1 (Figure 8J) in placental tissues from PETPIR-overexpressing mice, compared to the NC group. These molecular changes are consistent with the observed preeclampsia-like phenotype. These findings demonstrate that overexpression of PETPIR in pregnant mice recapitulates key features of preeclampsia, highlighting PETPIR as a critical mediator in the pathophysiology of preeclampsia.

## Discussion

Preeclampsia, a hypertensive disorder of pregnancy, remains a major cause of maternal and perinatal morbidity and mortality worldwide (2). While research has extensively focused on protein-coding genes and non-coding RNAs (29), the role of piRNAs, in preeclampsia has been largely overlooked (30). Addressing this gap, our study found PETPIR as a critical regulator in the pathophysiology of preeclampsia. We demonstrated that PETPIR is the most significantly upregulated piRNA in preeclamptic placentas, highlighting its potential as a biomarker. Functionally, PETPIR suppresses trophoblast proliferation, migration, and invasion—key processes for placental development and vascular remodeling—aligning with the impaired trophoblast function characteristic of preeclampsia (31, 32). Additionally, PETPIR modulates hypoxia-responsive pathways, suggesting its role in the hypoxic microenvironment of preeclamptic placentas (33). PETPIR enhances m6A RNA methylation by directly interacting with YTHDF2, a key m6A reader, thereby stabilizing downstream targets such as NDRG1, which were upregulated in both cellular and in vivo models. In pregnant mice, PETPIR overexpression recapitulated hallmark features of preeclampsia, including hypertension and fetal growth restriction, establishing a strong in vivo link between PETPIR dysregulation and preeclampsia pathogenesis.

PETPIR was identified as a significantly upregulated piRNA in preeclamptic placentas, with its conserved nature across species underscoring its functional importance in placental biology. This conservation suggests evolutionary pressure to preserve its regulatory roles in trophoblast function and placental development, aligning with the established roles of piRNAs in gene silencing, epigenetic regulation, and transposon suppression in germline cells (34–36). Functional analyses revealed that PETPIR overexpression inhibits trophoblast proliferation, migration, and invasion. Mechanistically, PETPIR induces cell cycle arrest by increasing the proportion of cells in the G0/G1 phase and reducing S-phase entry. PETPIR promotes apoptosis by upregulating pro-apoptotic markers, such as Bax and Casp3, and downregulating the anti-apoptotic marker Bcl-2. These findings align with the critical role of trophoblast proliferation and invasion in early placental development, where disruptions can lead to shallow trophoblast invasion and impaired spiral artery remodeling, hallmark features of preeclampsia (37). PETPIR further impairs trophoblast motility, as evidenced by significant reductions in cell migration and invasion. These functional deficits are likely to contribute to the insufficient remodeling of maternal spiral arteries characteristic of preeclampsia (38). Proper spiral artery remodeling, which involves the invasion of extravillous trophoblasts into the maternal decidua and the replacement of vascular endothelial and smooth muscle cells, ensures adequate blood flow to the developing fetus (39, 40). By suppressing these processes, PETPIR may directly contribute to the placental hypoperfusion and oxidative stress observed in preeclampsia. The identification of PETPIR as a piRNA involved in trophoblast biology provides novel insights into non-coding RNA-mediated regulation of placental development. While previous studies have focused on the roles of lncRNAs and miRNAs in trophoblast function and preeclampsia, our findings highlight a significant regulatory role for piRNAs. The ability of PETPIR to suppress trophoblast cell function positions it as a key molecular contributor to preeclampsia pathogenesis, offering a new direction for understanding this complex disorder.

RNA sequencing of PETPIR-overexpressing trophoblast cells revealed significant upregulation of genes involved in hypoxia-responsive pathways and HIF-1 signaling pathway. HIF-1, a master regulator of cellular responses to hypoxia, plays a critical role in placental development (41). Hypoxia, a hallmark of preeclampsia caused by impaired placental perfusion and shallow trophoblast invasion, results in reduced oxygen delivery to the placenta (42). PETPIR’s ability to enhance the expression of hypoxia-inducible genes highlights its role in modulating trophoblast responses to hypoxia and adapting to oxygen deprivation (43, 44)(45). Our findings also demonstrate that PETPIR enhances global m6A methylation and directly interacts with YTHDF2, a key m6A reader protein responsible for RNA stabilization and degradation (46). This interaction establishes a crucial mechanistic link by which PETPIR modulates the m6A epitranscriptome to regulate trophoblast cell function. Through YTHDF2-mediated stabilization of target transcripts, PETPIR acts as a critical post-transcriptional regulator in trophoblast biology and the pathophysiology of preeclampsia. By introducing a novel layer of post-transcriptional regulation, the PETPIR-YTHDF2 axis expands the role of piRNAs beyond their classical function in transposon silencing, connecting m6A modifications to essential trophoblast processes. In addition, our transcriptome-wide analysis of m6A methylation in preeclamptic placentas revealed extensive epitranscriptomic reprogramming, underscoring its critical role in the pathophysiology of preeclampsia. Hypermethylated and upregulated genes were enriched in pathways essential for placental function, including Tube Closure, HIF-1 signaling, Apical Junction Complex, and Tropomyosin Binding. These pathways are critical for maintaining placental vascular integrity, facilitating efficient oxygen and nutrient exchange, and supporting trophoblast function—all of which are severely compromised in preeclampsia (47). Among the hypermethylated and upregulated genes, IL6 and VEGFA emerged as central regulatory hubs within the gene interaction network, particularly in hypertension and cardiovascular diseases. These findings reflect their well-established roles in inflammation, angiogenesis, and hypoxia responses. The hypermethylation of these genes suggests that m6A-mediated regulation exacerbates hypoxia- and inflammation-driven pathologies, further contributing to the molecular dysfunction observed in preeclampsia.

Among PETPIR’s targets, NDRG1 emerged as a critical effector of the PETPIR/YTHDF2 axis. NDRG1, a well-characterized hypoxia-responsive gene (48–50), was significantly upregulated in PETPIR-overexpressing trophoblast cells and preeclamptic placentas, highlighting its role in mediating PETPIR’s pathogenic effects in preeclampsia. PETPIR enhances NDRG1 mRNA stability through YTHDF2. RIP assays confirmed that YTHDF2 directly binds to NDRG1 transcripts. Actinomycin D chase experiments further demonstrated that YTHDF2 overexpression slows NDRG1 mRNA degradation, thereby increasing transcript stability. These findings establish YTHDF2-mediated m6A methylation as a key regulatory mechanism through which PETPIR modulates NDRG1 expression. NDRG1, implicated in cellular stress responses, angiogenesis, and hypoxia adaptation, is known to be upregulated under hypoxic conditions to maintain cellular homeostasis (51, 52). In preeclampsia, its upregulation may represent a compensatory response to placental hypoxia. However, dysregulated or excessive NDRG1 expression could contribute to pathological features such as impaired trophoblast invasion and abnormal vascular remodeling, aligning with reports linking NDRG1 to preeclampsia-associated vascular dysfunction (53, 54). The identification of NDRG1 as a downstream effector of PETPIR provides new insights into the molecular pathogenesis of preeclampsia. By linking PETPIR-mediated m6A regulation to hypoxia-responsive gene expression, this study highlights how m6A alterations drive placental dysfunction. Moreover, NDRG1’s role in hypoxia and stress responses positions it as a potential therapeutic target for alleviating the effects of placental hypoxia in preeclampsia.

In vivo, overexpression of PETPIR in pregnant mice induced hallmark features of the disease, including elevated systolic blood pressure and fetal growth restriction. These phenotypes closely mimic the clinical manifestations of human preeclampsia, highlighting PETPIR’s translational relevance in disease pathophysiology. Histological analysis of uteroplacental tissues revealed significant disruptions in the lumen-to-wall ratio indicative of poor placental perfusion. Fetal growth restriction, evidenced by smaller placentas and lower fetal weights, underscores the downstream consequences of placental insufficiency driven by PETPIR overexpression. These in vivo results substantiate the functional importance of the PETPIR/YTHDF2/NDRG1 axis in preeclampsia pathogenesis. Future research should leverage this model to explore small-molecule inhibitors, RNA-based therapeutics, or other strategies aimed at modulating PETPIR expression or its downstream pathways. Additionally, the broader regulatory network of PETPIR warrants further investigation. The interaction of PETPIR with other RNA-binding proteins, non-coding RNAs, and upstream regulators remains an important area of exploration.

## Conclusion

In conclusion, our study provides foundational insights into the role of piwi-interacting RNA PETPIR in preeclampsia and establishes a framework for future investigations. By integrating molecular biology, epitranscriptomics, and preclinical models, this research paves the way for innovative strategies to manage preeclampsia and enhance maternal-fetal health outcomes.

## Data Availability

The data in support of the results are available from the corresponding author on reasonable request.

## Acknowledgements

NA

## Author contributions

Acquisition of data: Ying Zhao, Shuang Liang; Data analysis: Ying Zhao, Shuang Liang, Miao Guo, Shaoyuan Huang, Houzhi Yang, Shan-Shan Li and Xin Jin; Writing of the manuscript: Ying Zhao, Shan-Shan Li and Xin Jin; Review and/or revision of the manuscript: Xin Jin; Conception and design: Shan-Shan Li and Xin Jin; Study supervision: Xu Chen and Xin Jin.

